# Selective Elimination of HIV-1-Infected Macrophages by Targeted Activator of Cell Kill

**DOI:** 10.64898/2026.11.11.765927

**Authors:** Beatriz M Lahoz, Anouk AJ Hamers, Thaissa Cordeiro, Mario Stevenson

## Abstract

In the pursuit of an HIV cure, elimination of persistent HIV reservoirs requires strategies that promote immune-mediated or cytopathic clearance of infected cells. While many such approaches are being developed against infected CD4^+^ T cells, HIV-infected macrophages are comparatively resistant to cell death and to several killing mechanisms being explored for T-cell reservoirs, highlighting the need for strategies that specifically eliminate this reservoir. Targeted Activator of Cell Kill (TACK), developed by Merck Pharmaceuticals, are non-nucleoside reverse transcriptase inhibitor-derived molecules designed to selectively kill HIV-1-infected cells by inducing premature activation of the viral protease. Here, we evaluated the TACK compounds Pyr01 and it’s inactive analog Pyr02, in primary human monocyte-derived macrophages (MDMs). Pyr01 had minimal effects on uninfected MDM viability but potently targeted HIV-1-infected MDMs, with an EC₅₀ of approximately 0.36 μM. Pyr01 treatment produced a rapid and sustained reduction in extracellular virion-associated HIV-1 RNA and significantly decreased cell-associated viral RNA, whereas Pyr02 exhibited minimal activity. Immunofluorescence microscopy and live-cell imaging further demonstrated a marked loss of HIV-1-infected macrophages following Pyr01 treatment, consistent with selective elimination of infected cells. These findings identify Pyr01 as a specific mediator of HIV-infected macrophage death and establish premature viral protease activation as a strategy for overcoming the intrinsic resistance of macrophage reservoirs to clearance. Targeting this macrophage-specific barrier may complement approaches directed at CD4^+^ T-cell reservoirs and broaden the cellular reach of HIV cure strategies.

**IMPORTANCE:** A foundational principle in HIV cure strategies is the elimination of cells that harbor persistent virus. Although ART effectively suppresses HIV-1 replication, it prevents de novo viral replication but does not eliminate infected-cell reservoirs. Cure efforts have largely focused on CD4+ T-cell reservoirs, yet HIV-infected macrophages represent a distinct challenge because they are long-lived, can sustain productive infection, and resist killing mechanisms developed to eliminate infected CD4+ T cells. Here, we demonstrate that the TACK compound Pyr01 selectively kills HIV-1-infected primary human macrophages while largely preserving uninfected cells. These findings extend TACK activity beyond CD4+ T cells and identify premature activation of HIV protease as a strategy to overcome infected macrophage resistance to virus-mediated and host-mediated clearance. By targeting a cellular reservoir poorly addressed by existing cure approaches, this work expands the potential reach of TACK compounds and supports complementary strategies to eliminate HIV across distinct reservoir cell types.

## INTRODUCTION

Despite the success of combination antiretroviral therapy (ART) in durably suppressing HIV-1 replication, treatment is not curative because long-lived reservoir cells persist and can rapidly reignite viremia following ART interruption. Consequently, a fundamental objective of HIV cure strategies is not simply to halt virus replication, but to eliminate the cells that harbor it. Most cure efforts have focused on the well-characterized reservoir in CD4^+^ T cells and ultimately depend on viral cytopathic effects, immune-mediated killing, or therapeutically induced cell death to eliminate infected cells. Increasing evidence, however, indicates that cells of the myeloid lineage, particularly macrophages, can also harbor persistent HIV and may present a fundamentally different challenge to cure because of their intrinsic resistance to cell death.

Macrophages possess several properties that favor HIV persistence. They are long-lived, widely distributed throughout tissues and, in some anatomical compartments, maintained through local self-renewal. Unlike activated CD4^+^ T cells, macrophages are relatively resistant to HIV-induced cytopathic effects and can sustain productive infection for prolonged periods without undergoing cell death (1–3). Importantly, evidence from People with HIV (PWH) now supports the presence of persistent HIV within myeloid cells during suppressive ART. Ganor et al. demonstrated that urethral macrophages from ART-suppressed PWH contain integrated HIV DNA, viral RNA and protein and can release replication-competent virus following *ex vivo* stimulation (4). Veenhuis *et al*. subsequently identified inducible HIV in monocyte-derived macrophages from virologically suppressed PWH and detected intact proviruses in circulating monocytes from 40% of participants examined, with inducible reservoirs persisting longitudinally in some individuals (5).

Evidence that macrophage reservoirs can contribute to recrudescing viremia following ART interruption further underscores their potential relevance to cure. Andrade *et al*. identified macrophage-tropic HIV-1 variants among rebound viruses from individuals undergoing analytical treatment interruption (ATI) (6). Some macrophage-tropic virions carried the macrophage-associated surface marker CD14, consistent with their production by macrophages, and phylogenetic analyses indicated that these viral lineages existed before treatment interruption. Thus, macrophage-lineage cells present during suppressive ART may contribute directly to the pool of virus that emerges when therapy is withdrawn.

The central nervous system provides additional evidence for a durable tissue macrophage-lineage reservoir. Using brain tissue obtained by rapid autopsy from ART-treated PWH, Tang *et al.* isolated highly purified brain myeloid cells that predominantly expressed the microglial marker TMEM119 and contained total and integrated HIV DNA together with low levels of viral RNA (7). Furthermore, HIV could be induced from parietal-cortex microglia and recovered virus productively infected both microglia and PBMCs. Brain-derived virus was macrophage tropic and genetically distinct from peripheral viral populations, providing direct evidence that brain-resident microglia can harbor inducible, replication-competent HIV despite durable ART. Collectively, these human studies provide complementary evidence that macrophage-lineage cells can harbor persistent HIV during ART and may contribute to viral rebound if ART suppression is lifted.

The importance of the macrophage reservoir is amplified by a second challenge: HIV/SIV-infected macrophages are unusually resistant to elimination. In addition to tolerating productive viral replication without succumbing to viral cytopathicity (1–3), infected macrophages resist immune effector mechanisms upon which many HIV cure strategies depend. Vojnov *et al*. first demonstrated that freshly isolated SIV-specific CD8^+^ T cells that efficiently eliminated infected CD4^+^ T cells and suppressed viral replication failed to e eliminate SIV-infected macrophages or suppress virus production from these cells (8). Subsequent studies by Rainho *et al.* demonstrated that this resistance persisted when Nef-mediated MHC-I downregulation was disrupted, indicating that macrophage resistance to CTL killing cannot be explained simply by impaired antigen recognition and instead reflects intrinsic properties of the infected macrophage (9).

These observations were subsequently extended to HIV-infected human macrophages. Clayton *et al*. demonstrated that HIV-infected macrophages were substantially more resistant than infected CD4^+^ T cells to CTL-mediated killing, resulting in inefficient suppression of HIV replication (10). Macrophage killing was delayed and dependent upon granzyme B and caspase-3, in contrast to the rapid killing observed in infected CD4^+^ T cells. Resistance also extends to innate immune effectors: HIV-infected macrophages resist efficient NK cell-mediated cytotoxicity, with NK-cell engagement favoring inflammatory cytokine responses rather than efficient cytolytic activity (11). Together, these studies identify a fundamental challenge for HIV cure: macrophages can harbor persistent, biologically competent HIV yet resist viral cytopathicity as well as adaptive and innate immune mechanisms that would otherwise eliminate infected cells.

Our previous work demonstrated that this resistance is therapeutically tractable. HIV-infected macrophages can be sensitized to cell death through manipulation of host survival pathways, including CSF1R antagonism in combination with TRAIL (12). These findings provide proof-of-concept that macrophage survival mechanisms can be disrupted but also underscore the potential limitations of approaches requiring multiple agents or manipulation of broadly expressed host pathways. An alternative strategy would be to directly couple expression of HIV proteins to death of the infected macrophage, thereby exploiting the virus itself to selectively eliminate the cell in which it resides and reducing dependence on immune effector mechanisms to which macrophages are resistant.

Agents such as Targeted Activator of Cell Kill (TACK), developed by Merck Pharmaceuticals, offer such an approach. TACK compounds are non-nucleoside reverse transcriptase inhibitor (NNRTI)-derived molecules engineered to selectively eliminate HIV-expressing cells (13). Conventional NNRTIs inhibit reverse transcription through allosteric binding to the p66 subunit of HIV reverse transcriptase (RT) (14). TACK retains this activity but additionally, binds the RT domain within monomeric Gag-Pol to accelerate its dimerization, triggering premature intracellular activation of HIV protease. This aberrant protease activity results in cleavage of viral and cellular proteins and activation of cell-death pathways, including the CARD8 inflammasome and pyroptotic cell death (13–17). Balibar et al. developed analogs of the NNRTI Doravirine, with markedly increased ability to promote death of HIV-expressing cells at pharmacologically relevant concentrations, including infected CD4^+^ T cells obtained from PWH (13). Because TACK activity couples expression of viral proteins directly to death of the infected cell, it offers a particularly attractive strategy for eliminating macrophages that resist conventional viral and immune-mediated cytotoxicity.

Among compounds identified in the TACK screen of Balibar et al. (13), Pyrimidone 01 (Pyr01) exhibited potent cytotoxic activity, whereas the structurally related Pyr02 showed minimal cytotoxic activity. This functional distinction provides an opportunity to determine whether TACK-mediated activation of an intracellular viral death pathway can overcome the intrinsic resistance of primary macrophages to elimination. Although TACK compounds have demonstrated potent activity in HIV-infected T-cell systems, their capacity to selectively eliminate HIV-infected primary human macrophages remains unknown.

Here, we tested the hypothesis that premature activation of HIV protease by TACK compounds can overcome the intrinsic resistance of infected macrophages to cell death. We compared Pyr01 and Pyr02 in HIV-1-infected primary human monocyte-derived macrophages and demonstrate that Pyr01 selectively eliminates infected macrophages while largely preserving uninfected cells. These findings extend TACK-mediated killing to a reservoir cell type that is poorly eliminated by viral cytopathicity and host immune effectors and support targeted activation of an intracellular viral death pathway as a complementary strategy for eliminating macrophage reservoirs in HIV cure efforts.

## RESULTS

### TACK compounds exhibit minimal cytotoxicity in human macrophages

To determine whether TACK compounds exhibit nonspecific cytotoxicity, primary human monocyte-derived macrophages (MDMs) were treated with increasing concentrations of Pyr01, Pyr02, efavirenz (EFV; a conventional NNRTI), or staurosporine (STA; a potent inducer of apoptosis), and viability was assessed by MTT assay (Fig. 1A). Cell viability was normalized to untreated MDMs (100%). Pyr01, Pyr02, and EFV had minimal effects on macrophage viability at concentrations up to 100 μM, with reduced viability observed only at 100 μM. In contrast, STA produced a marked, concentration-dependent loss of viability, confirming assay sensitivity (18, 19) (Fig. 1B). DMSO vehicle controls, matched to the volumes used at each compound concentration, had no detectable effect on viability. Thus, Pyr01 and Pyr02 exhibit minimal cytotoxicity toward uninfected primary macrophages at concentrations previously shown to be active in CD4^+^ T cells (13), supporting evaluation of their activity against HIV-infected macrophages.

**Figure 1:**
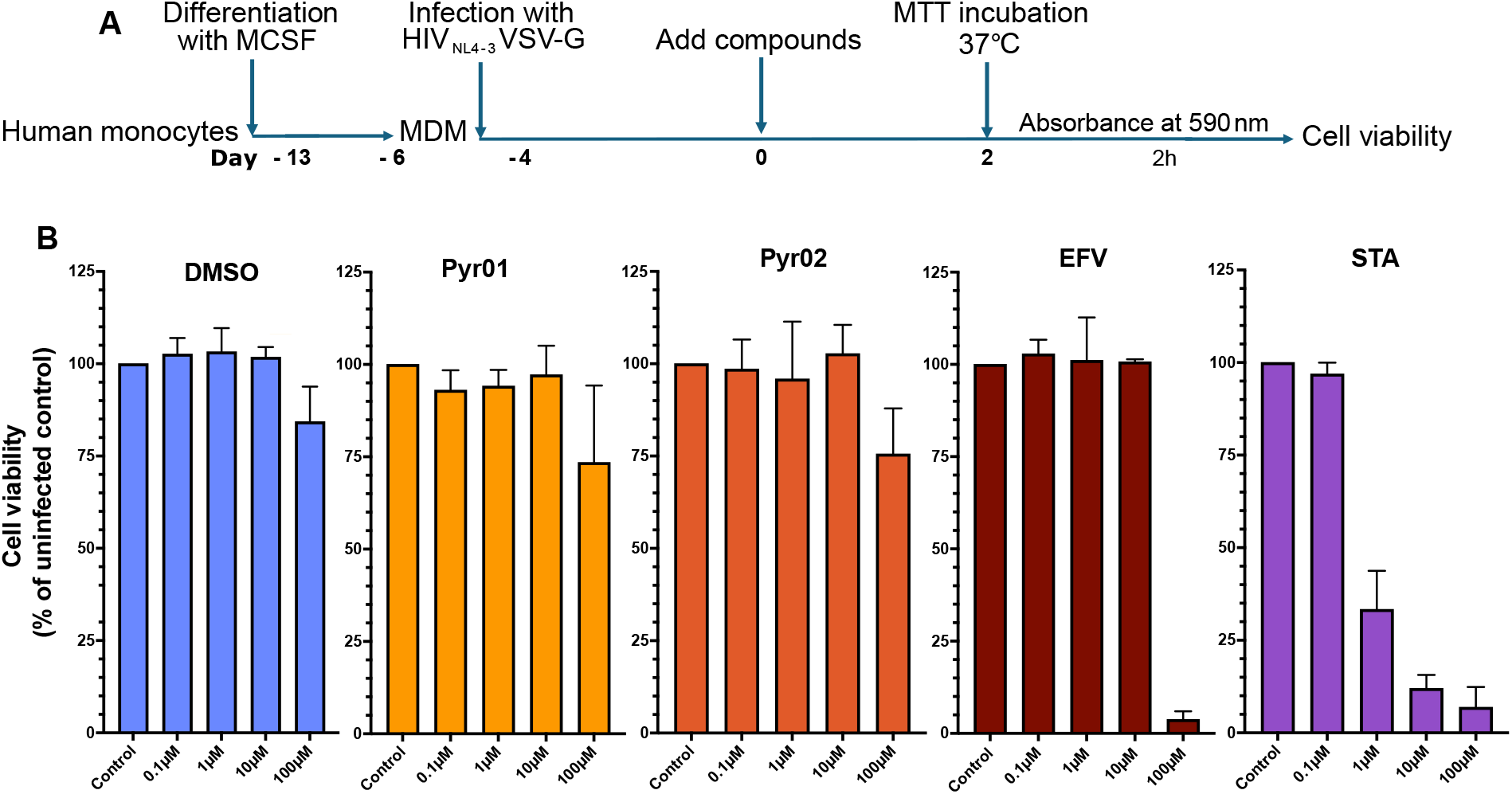
Effect of Pyr01, Pyr02, efavirenz, and staurosporine on primary human macrophage viability. (A) Primary human monocytes from three independent healthy donors were differentiated into MDM using M-CSF. Cells were treated with increasing concentrations (0.1, 1, 10, or 100 μM) of Pyr01, Pyr02, efavirenz (EFV), or staurosporine (STA). Corresponding quantities of DMSO were taken along as vehicle control. (B) Cell viability was determined by MTT assay after 2 days of treatment and expressed as a percentage of the untreated control, which was set to 100%. Data are presented as mean ± SD.

### Dose-response analysis identifies Pyr01 as a potent inhibitor of HIV-1 infection in primary human macrophages

We next determined whether Pyr01, previously shown to promote killing of HIV-1-infected CD4^+^ T cells, could similarly reduce infected primary human macrophages. MDMs were infected overnight with VSV-G-pseudotyped HIV-1NL4-3 to restrict infection to a single round. Four days after infection, cells were treated with serial dilutions of Pyr01 or Pyr02, and the frequency of HIV-1 Gag^+^ MDMs was quantified by immunofluorescence two days later (Fig. 2A). Pyr01 produced a dose-dependent reduction in Gag^+^ macrophages across four independent donors, whereas Pyr02 had little or no effect at the concentrations tested (Fig. 2B). Nonlinear regression yielded an EC₅₀ of 0.36 μM (95% CI, 0.22–0.60 μM) for Pyr01 (Fig. 2C), while the absence of a concentration-dependent response to Pyr02 precluded reliable EC₅₀ determination. These results demonstrate potent, dose-dependent activity of Pyr01 against HIV-1-infected primary macrophages and provide initial evidence that TACK activity extends beyond CD4^+^ T cells to macrophages.

**Figure 2:**
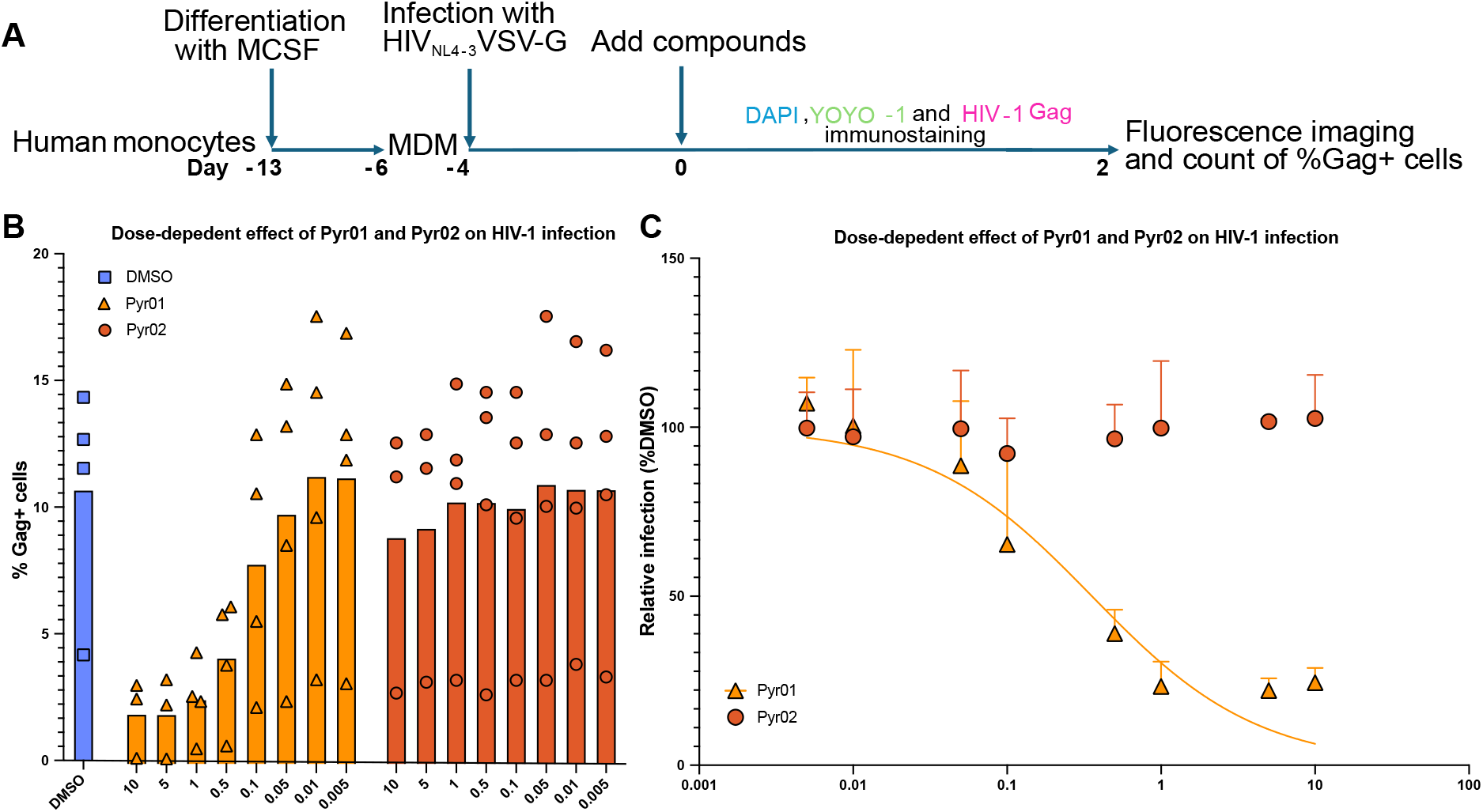
Dose-dependent inhibition of HIV-1 infection by Pyr01 in primary human macrophages. (A) Human monocytes were differentiated into MDM, infected with VSV-G-pseudotyped HIV-1_NL4-3_, and treated with serial dilutions of Pyr01 or Pyr02 (ranging from 5 nM to 10 μM). Two days after treatment, cells were fixed and stained with DAPI, YOYO-1, and anti-HIV-1 Gag antibody. (B) Infection was quantified by fluorescence microscopy as the percentage of Gag^+^ cells. Data represent three technical replicates from each of four independent donors. Data are presented as mean ± SEM. (C) Nonlinear regression analysis was performed using a variable-slope dose-response model. Infection data was normalized to the corresponding DMSO-treated control for each donor. Pyr01 exhibited a concentration-dependent reduction in HIV-1 infection with an estimated EC₅₀ of 0.36 μM, whereas Pyr02 did not produce a measurable dose-dependent inhibitory effect across the concentrations tested. Data are presented as geometric mean ± SD.

### Pyr01 inhibits productive HIV-1 infection of human macrophages

To determine the effect of TACK treatment on viral production, extracellular HIV-1 RNA was quantified daily in supernatants from infected MDM cultures for four days following treatment (Fig. 3A). Individual profiles from four donors, shown as the mean ± SD of technical replicates, are presented in Figure 3B. To account for donor-to-donor variation in infection, values were normalized to the corresponding DMSO vehicle control and combined for analysis (Fig. 3C). Pyr01 produced a rapid and sustained reduction in extracellular HIV-1 RNA, with levels approaching the limit of detection throughout the treatment period. Pyr02 and EFV also reduced extracellular viral RNA but were consistently less effective than Pyr01. Pairwise comparisons confirmed significantly greater activity of Pyr01 compared with Pyr02 at multiple time points (Fig. 3D).

**Figure 3:**
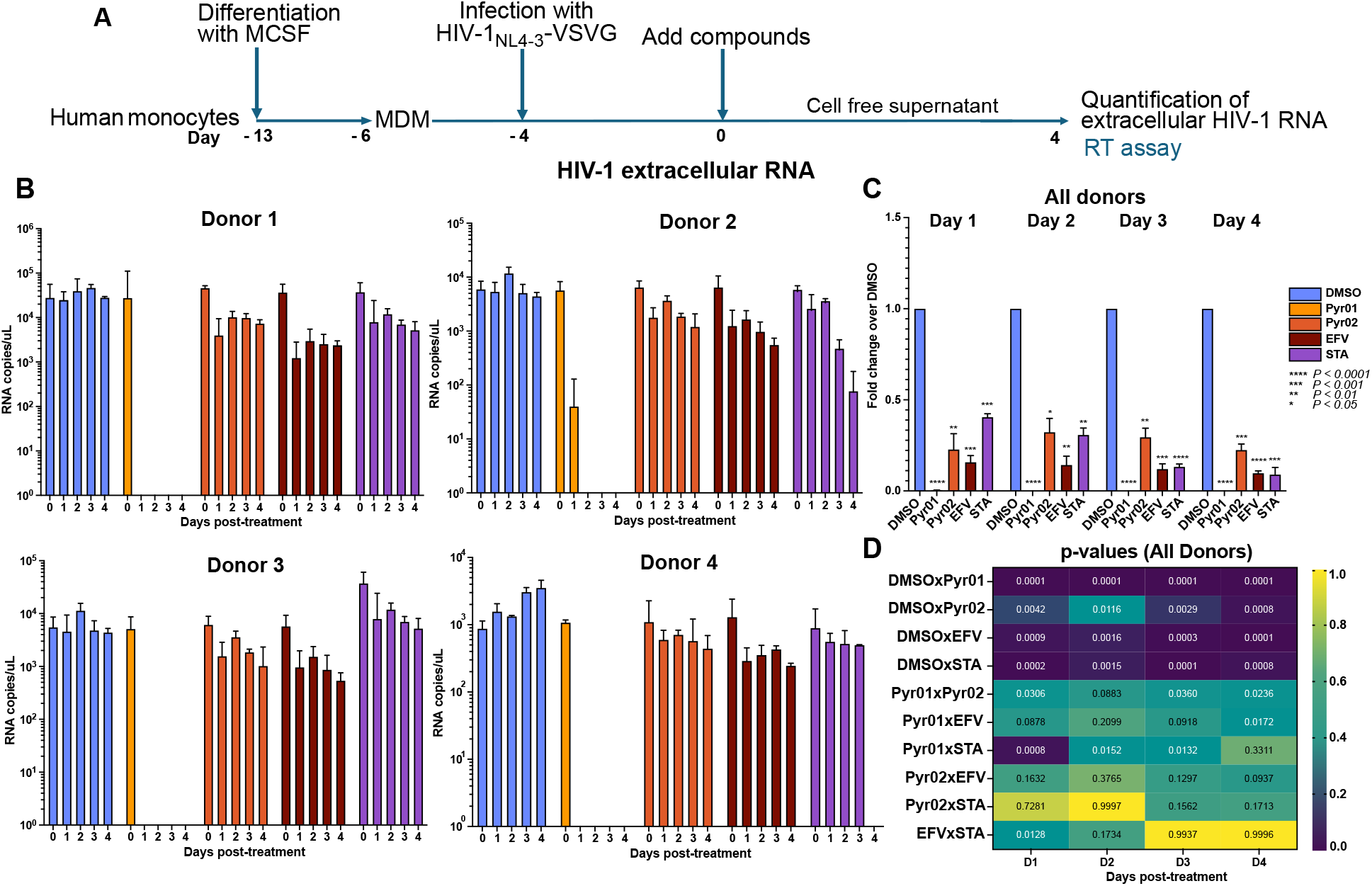
Pyr01 eliminates the number of virions in supernatants from HIV-1-infected primary human macrophages. (A) Human monocytes from four independent healthy donors were differentiated into MDM and infected with VSV-G-pseudotyped HIV-1_NL4-3_. Four days after infection, cells were treated with either DMSO, Pyr01, Pyr02, efavirenz (EFV), or staurosporine (STA), with Day 0 indicating the day of compound addition. (B) Culture supernatants were collected daily (Days 1–4), and virion-associated HIV-1 RNA was quantified by direct RT-qPCR. Data from multiple technical replicates for each individual donor are presented as mean ± SD. (C) Combined normalized data from all 4 donors. To account for baseline biological variability shown in panel B, data were normalized within each donor and are expressed as mean fold change relative to the respective DMSO vehicle control ± SEM. Statistical significance was determined using an ordinary two-way ANOVA followed by Tukey’s multiple-comparisons test. Statistical significance of treatments versus DMSO control at each time point: **** *p* < 0.0001; *** *p* < 0.001; ** *p* < 0.01; * *p* < 0.05. (D) Heat map displaying *p*-values for all possible comparisons among treatment groups at days 1-4 post-treatment, generated from the normalized fold-over-DMSO data shown in panel C. Cell colors and text indicate exact *p*-values.

We next determined whether the reduction in extracellular virus was accompanied by a decrease in cell-associated HIV-1 RNA. Infected MDMs were harvested daily following treatment, and intracellular viral RNA was quantified by RT-qPCR (Fig. 4A). Pyr01 consistently reduced cell-associated HIV-1 RNA across individual donors throughout the treatment period (Fig. 4B). To account for differences in baseline infection, values were normalized to the corresponding DMSO control for each donor (Fig. 4C). Pyr01 significantly reduced cell-associated HIV-1 RNA at every time point examined, whereas Pyr02 had no significant effect, and EFV showed only a trend toward reduction. Pairwise analyses further demonstrated significantly greater reductions with Pyr01 than Pyr02 at multiple time points, with EFV producing an intermediate effect (Fig. 4D).

**Figure 4:**
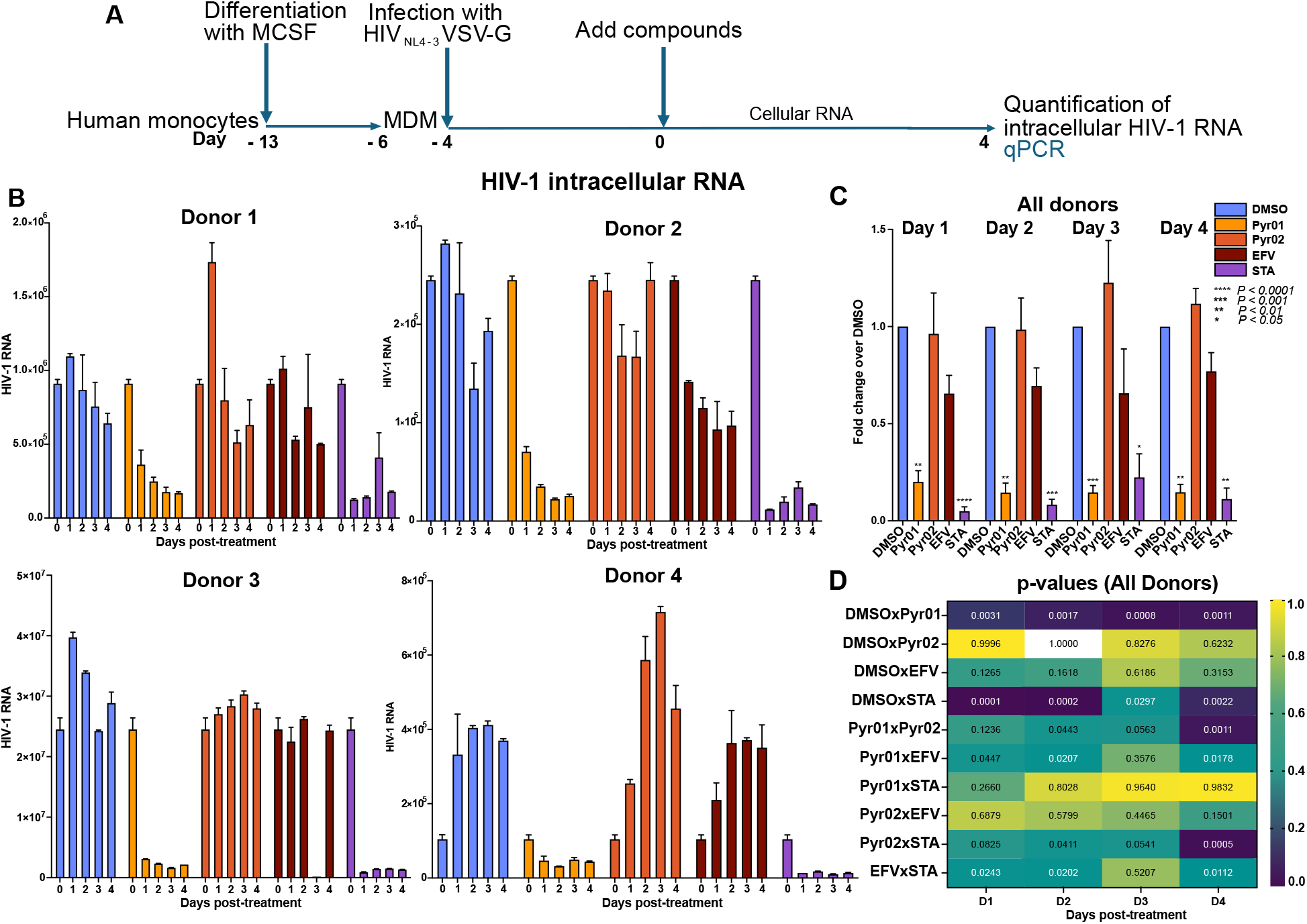
Pyr01 reduces cell-associated HIV-1 RNA in primary human macrophages. (A) Monocytes from four independent healthy donors were differentiated into MDM and infected with VSV-G-pseudotyped HIV-1_NL4-3_. Four days after infection, cells were treated with DMSO, Pyr01, Pyr02, efavirenz (EFV), or staurosporine (STA), with Day 0 indicating the day of compound addition. Cell lysates were harvested daily (Days 1–4), followed by RNA extraction. (B) Cell-associated HIV-1 RNA was quantified by RT-qPCR. Data from multiple technical replicates for each individual donor are presented as mean ± SD. (C) Combined normalized data from all 4 donors. To account for baseline biological variability shown in panel B, data were normalized within each donor and are expressed as mean fold change relative to the respective DMSO vehicle control ± SEM. Statistical significance was determined using an ordinary two-way ANOVA followed by Tukey’s multiple-comparisons test. Statistical significance of treatments versus DMSO control at each time point: **** *p* < 0.0001; *** *p* < 0.001; ** *p* < 0.01; * *p* < 0.05. (D) Heat map displaying *p*-values for all possible comparisons among treatment groups at days 1-4 post-treatment, generated from the normalized fold-over-DMSO data shown in panel C. Cell colors and text indicate exact *p*-values.

Together, these findings demonstrate that Pyr01 produces a rapid and sustained reduction in both extracellular and cell-associated HIV-1 RNA in primary human macrophages. The greater activity of Pyr01 relative to the structurally related, TACK-inactive Pyr02 supports a TACK-dependent effect rather than antiviral activity attributable solely to the NNRTI scaffold.

### Pyr01 selectively eliminates HIV-1-infected primary human macrophages

To directly assess the effect of TACK compounds on HIV-1-infected macrophages, MDMs infected with VSV-G-pseudotyped HIV-1NL4-3 were treated for two days and analyzed by immunofluorescence microscopy. Nuclei were identified by DAPI, infected cells by HIV-1 Gag staining, and nonviable cells by YOYO-1 staining (Fig. 5A). Pyr01 treatment produced a marked reduction in Gag^+^ macrophages compared with vehicle control (p=0.0005). Importantly, this reduction was not accompanied by a substantial increase in YOYO-1 staining, indicating preservation of overall macrophage viability and supporting selective elimination of HIV-1-infected cells. In contrast, Pyr02 and EFV did not significantly reduce Gag^+^ cells and produced minimal YOYO-1 staining. As expected, staurosporine (STA) caused extensive nonspecific cell death, reflected by a marked loss of macrophages and widespread YOYO-1 positivity irrespective of Gag expression.

**Figure 5:**
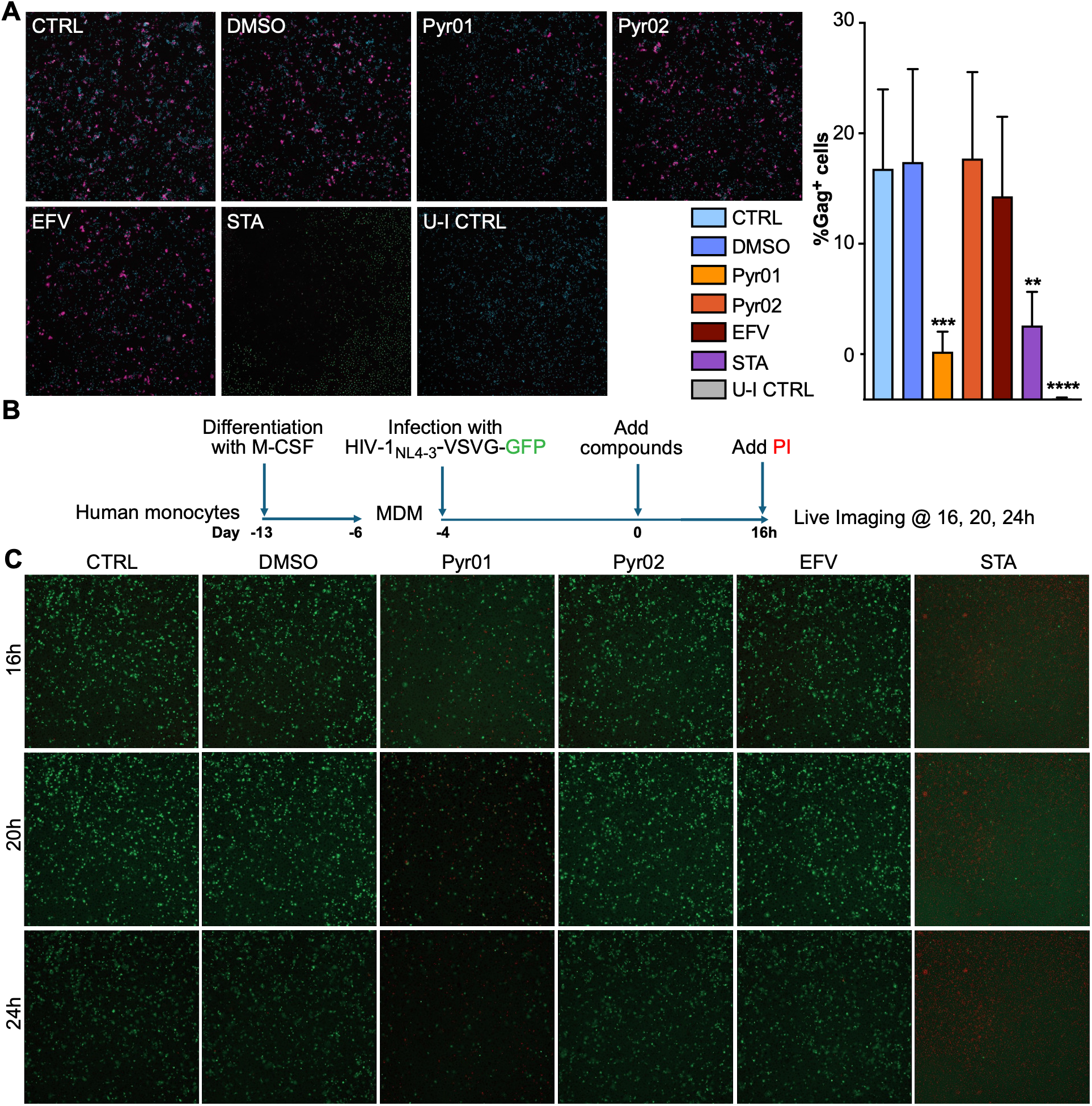
Pyr01 strongly diminishes the number of HIV-1-infected primary human macrophages. (A) Monocytes from five independent healthy donors were differentiated into MDM and infected with VSV-G-pseudotyped HIV-1_NL4-3_. Four days after infection, cells were treated with DMSO (= vehicle), Pyr01, Pyr02, efavirenz (EFV), or staurosporine (STA). Two days after treatment, cells were fixed and stained with DAPI (nuclei, blue), YOYO-1 (dead cells, green), and anti-HIV-1 Gag antibody (magenta). Center well images were acquired for visualization, and the entire well was imaged to quantify the percentage of Gag^+^ cells. Data are presented as mean ± SD. Statistical significance was determined using an ordinary one-way ANOVA followed by Dunnett’s multiple-comparisons test. Statistical significance of each treatment versus DMSO control: **** *p* < 0.0001; *** *p* < 0.001; ** *p* < 0.01. CTRL = infected untreated control; U-I CTRL = uninfected untreated control. (B) Monocytes from one healthy donor were differentiated into MDM and infected with VSV-G-pseudotyped HIV-1_NL4-3_ GFP reporter virus. Four days after infection, cells were treated with DMSO, Pyr01, Pyr02, EFV, or STA. 16hrs after treatment, PI was added and live imaging was performed at 16, 20, and 24 hours. (C) Representative center well images of each treatment at each timepoint are presented. Dead cells are indicated with PI in red and HIV^+^ cells with GFP in green.

We next used live-cell imaging to examine the kinetics and selectivity of Pyr01 activity. MDMs infected with a VSV-G-pseudotyped HIV-1NL4-3 GFP reporter virus were treated with Pyr01, Pyr02, EFV, or DMSO and monitored longitudinally, with propidium iodide (PI) used to identify nonviable cells (Fig. 5B). Pyr01 produced a progressive loss of GFP^+^ macrophages over time (Fig. 5C), whereas Pyr02 and EFV had little effect on the number of GFP^+^ cells. Importantly, loss of GFP^+^ cells following Pyr01 treatment occurred without widespread PI positivity, consistent with selective elimination of infected macrophages rather than nonspecific cytotoxicity. In contrast, STA caused rapid loss of GFP signal accompanied by extensive PI positivity, while DMSO-treated cultures maintained GFP^+^ macrophages throughout the observation period.

Together, several observations support selective elimination of infected cells, rather than nonspecific macrophage toxicity, as an important component of Pyr01 activity. First, Pyr01 exhibited minimal cytotoxicity toward uninfected macrophages at concentrations that potently reduced infection. Second, fixed-cell and longitudinal live-cell imaging demonstrated a progressive loss of HIV Gag^+^ or GFP^+^ macrophages without widespread death of the surrounding macrophage population. Third, Pyr02, despite its structural similarity to Pyr01, failed to produce a comparable reduction in infected cells. Finally, EFV reduced viral replication but did not substantially reduce the infected cell population. These distinctions are consistent with the proposed TACK mechanism, in which binding to the reverse transcriptase domain of Gag-Pol promotes premature RT dimerization and intracellular HIV protease activation, ultimately triggering CARD8 inflammasome-mediated pyroptotic death (13, 15–17). Thus, unlike conventional antiretroviral drugs that suppress virus production while leaving the infected cell intact, TACK compounds have the potential to convert expression of an essential viral protein complex into a signal for elimination of the cell producing it.

## DISCUSSION

A central challenge for HIV cure is not simply to expose persistent virus, but to eliminate the cells that harbor it. This challenge may be particularly important for macrophages, which can sustain HIV infection while resisting viral cytopathicity and immune-mediated clearance. A particularly important implication of our findings is that TACK compounds can promote the elimination of HIV-infected macrophages, a cell population that is unusually resistant to both virus-induced and host-mediated cell death. Across independent donors, Pyr01 produced rapid and sustained reductions in extracellular HIV-1 RNA, cell-associated viral RNA, and the frequency of infected macrophages. Importantly, these effects occurred with minimal toxicity to uninfected macrophages. In contrast, the structurally related but TACK-inactive compound Pyr02 exhibited little activity, while efavirenz (EFV) suppressed viral replication without producing the pronounced loss of infected cells observed with Pyr01.

Unlike activated CD4^+^ T cells, macrophages can sustain productive HIV infection for prolonged periods without undergoing rapid cytopathic death. Studies of macrophages and microglia demonstrated that HIV-expressing cells are relatively resistant to apoptosis, supporting the concept that survival during productive infection may contribute to persistence of the myeloid reservoir (20). More recent studies similarly demonstrated that HIV-infected macrophages resist the cytopathic effects of acute infection and upregulate multiple pro-survival factors, including XIAP, cIAP1, BCL-2, and BCL-XL (21). Thus, the ability of Pyr01 to induce loss of infected macrophages is significant because it appears to overcome a fundamental biological property that distinguishes macrophages from the more cytopathicity-susceptible CD4^+^ T-cell reservoir.

HIV infection can further reinforce macrophage survival by actively engaging host anti-apoptotic pathways. We previously demonstrated that HIV-1 Env induces macrophage colony-stimulating factor (M-CSF), resulting in downregulation of the death receptor TRAIL-R1/DR4 and increased expression of the anti-apoptotic proteins Bfl-1 and Mcl-1. Disruption of this pathway partially restored susceptibility of infected macrophages to TRAIL-mediated apoptosis (3). Subsequent studies demonstrated that HIV-induced activation of CSF1R contributes to this phenotype and that pharmacologic CSF1R antagonism can sensitize infected macrophages to TRAIL-mediated killing (12). Together, these observations indicate that persistence of infected macrophages is not simply a consequence of their intrinsic longevity; HIV can actively exploit macrophage survival pathways to sustain the infected cell in the face of host immune clearance processes.

Resistance also extends to adaptive immune-mediated clearance. In the SIV model, freshly isolated virus-specific CD8^+^ T cells that efficiently eliminated infected CD4^+^ T cells were markedly less effective at suppressing replication in or eliminating infected macrophages (8). Importantly, this resistance could not be explained simply by Nef-mediated downregulation of MHC-I. Disruption of Nef functions that impair MHC-I expression enhanced elimination of infected CD4^+^ T cells but failed to restore rapid CD8^+^ T-cell-mediated killing of infected macrophages (9). These observations were subsequently extended to HIV, where direct comparison of infected macrophages and CD4^+^ T cells demonstrated substantially less efficient CTL-mediated killing of macrophages. Macrophage killing was delayed and dependent on granzyme B and caspase-3, whereas killing of infected CD4^+^ T cells was considerably more rapid (10). Collectively, these studies suggest that resistance occurs downstream of immune recognition and reflects, at least in part, fundamental differences in how macrophages respond to cytotoxic signals.

HIV-infected macrophages are similarly resistant to innate immune effector mechanisms. NK cells efficiently recognize and eliminate HIV-infected CD4^+^ T cells, including through antibody-dependent cellular cytotoxicity (ADCC). In contrast, HIV-infected macrophages exhibit markedly reduced susceptibility to NK-cell killing and ADCC despite accessibility of HIV Env on the target cell. NK-cell interactions with infected macrophages instead favor inflammatory cytokine production over cytolytic activity (11). This distinction has important implications for HIV cure approaches that rely on antibodies or endogenous immune effectors: recognition of an infected macrophage does not necessarily translate into its efficient elimination.

Taken together, these studies identify a convergence of mechanisms that favor survival of the HIV-infected macrophage: resistance to direct viral cytopathicity, HIV-induced activation of cellular survival pathways, resistance to death-receptor-mediated apoptosis, inefficient CD8^+^ T-cell cytotoxicity, and impaired NK-cell-mediated killing and ADCC. This convergence may help explain how macrophages can sustain productive infection and persist as long-lived sources of virus despite immune recognition. Importantly, it also suggests that strategies optimized for elimination of infected CD4^+^ T cells cannot be assumed to efficiently eliminate macrophage reservoirs.

Our findings position TACK compounds as a mechanistically distinct approach to this problem. Rather than relying on an external death ligand or immune effector to overcome macrophage survival pathways, TACK compounds exploit a viral process within the infected cell. Pyr01 promotes premature activation of HIV protease through its interaction with Gag-Pol, thereby converting expression of viral proteins into a trigger for infected-cell death. The selective loss of Gag^+^ and GFP^+^ macrophages observed in our experiments, together with preservation of the surrounding uninfected macrophage population, suggests that this mechanism can circumvent at least some of the intrinsic and virus-induced barriers that normally protect HIV-infected macrophages from elimination.

This distinction may be particularly important for cure strategies. Approaches based on latency reversal, therapeutic vaccination, bNAbs, or other immune interventions ultimately depend on effective clearance of cells in which HIV expression has been induced (22–25). For macrophages, however, viral reactivation or immune recognition alone may be insufficient if the infected cell remains resistant to the resulting cytotoxic signal. TACK compounds offer the possibility of coupling viral expression directly to cell death, potentially bridging the gap between reservoir reactivation and reservoir elimination. The ability of Pyr01 to selectively eliminate HIV-expressing macrophages therefore extends the significance of TACK beyond its previously demonstrated activity in CD4^+^ T cells and suggests a complementary strategy for targeting a cellular reservoir that is intrinsically difficult to clear.

These findings also provide a rationale for evaluating TACK compounds in more physiologically relevant models of macrophage persistence, including long-lived tissue macrophages and microglia, and in combination with interventions that induce viral expression. Because TACK activity requires expression of the Gag-Pol substrate, deeply latent macrophages may remain insensitive until viral transcription and translation are induced. Conversely, once HIV expression occurs, TACK-mediated premature protease activation could provide an intracellular mechanism for eliminating a cell that might otherwise survive viral reactivation and evade immune-mediated killing. Thus, the particular significance of TACK for HIV cure may lie not simply in its ability to inhibit HIV replication, but in its capacity to exploit viral protein expression to overcome the exceptional resistance of HIV-infected macrophages to clearance.

## MATERIALS AND METHODS

### Cell Culture

All cell cultures were maintained under standard culture conditions (37°C, 5% CO₂) with media replenished as needed. HEK293T cells were obtained from ATCC and maintained in Dulbecco’s Modified Eagle Medium (DMEM; Gibco) supplemented with 10% heat-inactivated fetal bovine serum (Cytivia), 100 units/mL penicillin (Gibco), 100 μg/mL streptomycin (Gibco), and 2 mM L-glutamine (Gibco). Primary human monocyte-derived macrophages (MDMs) were generated from elutriated peripheral blood obtained from healthy donors through The Elutriation Core Facility at the University of Nebraska Medical Center. Total monocytes were isolated by negative selection using a commercially available kit (StemCell Technologies) according to the manufacturer’s instructions. Monocytes were differentiated into macrophages in differentiation medium consisting of DMEM supplemented with 10% heat-inactivated human serum (Sera Care Life Sciences), 100 units/mL penicillin, 100 μg/mL streptomycin, 2 mM L-glutamine and 13 ng/mL human recombinant monocyte colony-stimulating factor (rhM-CSF) (R&D Systems). After 7 days, MDMs were seeded for different experiments in D10 media (= differentiation media without rhM-CSF). Cells were allowed to rest and adhere for three days prior to infection and/or treatments to ensure recovery and stabilization.

### Preparation of viral stocks

Viral stocks were generated in HEK293T cells by co-transfection with the fuGENE HD Transfection Reagent (Promega), in opti-MEM (Sigma), and plasmids encoding the HIV-1 molecular clone NL4-3 or NL4-3 GFP, together with the vesicular stomatitis virus glycoprotein (VSV-G), using a 7:1 ratio. Following transfection, culture media was replaced after 18 hours to remove transfection reagents and improve cell viability. After an additional overnight incubation, virus-containing supernatants were harvested, and cellular debris was removed by passing through a 0.45 μm filter (CELLTREAT). Viral stocks were aliquoted and stored at −80°C for single use. Viral preparations were quantified by RT-qPCR prior to infection to ensure consistent input across experimental conditions.

### Infection and Treatment

Four days after infection, MDM were treated with 1 μM Pyr01, Pyr02, or efavirenz (EFV) unless mentioned otherwise. Pyr01 and Pyr02 were provided as powders by Merck and dissolved in dimethyl sulfoxide (DMSO; Sigma-Aldrich). EFV (Sigma-Aldrich), a clinically approved NNRTI, was included as a comparator to distinguish classical reverse transcriptase inhibition from TACK-mediated effects. Staurosporine (STA; Sigma-Aldrich) was used at 25 μM as a positive control for cytotoxicity, and DMSO served as the vehicle control. The final DMSO concentration in all treatment conditions was 0.01%. MDM were infected with VSV-G–pseudotyped HIV-1NL4-3, which promotes efficient and synchronous infection independent of CD4 and CCR5 expression, at 6.25 million RNA copies per 50,000 cells. Cells were exposed to the viral inoculum overnight to allow productive infection, after which the inoculum was removed, and cells were washed to eliminate unbound virus. Infected cultures were then maintained in D10 medium and monitored longitudinally for viral production and treatment effects.

### Evaluation of drug toxicity

50,000 MDM were seeded into 96-well plates and treated with serial dilutions of Pyr01, Pyr02, EFV, or STA (0.1 - 100 μM) under conditions matching those used in the infection experiments. DMSO was used as vehicle control. 2 days later, cell viability was assessed using an MTT assay as described previously (26). In short, cells were washed with warm DPBS (Gibco) and incubated with 200 μL of 0.5 mg/mL of the tetrazolium dye MTT (Sigma-Aldrich) in D10 for 1 h at 37°C. The MTT solution was removed; the cells were lysed with 300 μL isopropanol and absorbance was read at 590 nm (Fig. 1A). To correct for background absorbance, the values of 4 cell-free wells were averaged and subtracted from all raw experimental data. Cell viability was calculated as a percentage of the untreated control by dividing the absorbance of the treated cells with the absorbance of the control cells and then multiplying that value by 100.

### TACK dose-response

50,000 MDM were seeded into 96-well plates, infected VSV-G pseudotyped HIV-1NL4-3 and treated with a serial dilution of Pyr01 and Pyr02 across a broad concentration range (5 nM to 10 μM). After 2 days, cells were stained for viability, HIV-1 Gag, and nuclei as described below under “Fluorescence imaging” and in Figure 2A. Infection was quantified by high-content fluorescence imaging based on the percentage of Gag-positive cells. For each biological donor, infection levels were normalized to the corresponding DMSO-treated control (defined as 100% infection). Concentration-response curves were generated by nonlinear regression using ([Inhibitor] vs. normalized response) in GraphPad Prism (version 11.0.2), and half-maximal effective concentrations (EC₅₀) with 95% confidence intervals were calculated from the fitted curves.

### Analysis of extracellular HIV-1 RNA

Supernatants were collected daily following peak infection to monitor viral production over time. Virion-associated RNA was quantified by direct RT-qPCR targeting the HIV-1 gag region using primers adapted from a previously described assay (27). Direct RNA amplification was performed using Volcano 2G DNA Polymerase (Medix Biochemica; Cat. No. 8100M) as previously described by Sharkey et al. (28). Briefly, supernatants were diluted 1:20 in nuclease-free water to reduce PCR inhibition, and 2 μL of diluted sample was added to a 20-μL reaction containing 5× Volcano buffer, dNTPs, anti-Taq antibody (Invitrogen), LA9115/f primer (5′-TGCTTTTTGCCTGTACTGGGTCT-3′), 8T22 primer (5′-TTTTTTTTTTTTTTTTTTTTTTTGAAGCAC-3′), 2nr4nrB probe (5′-CCACTGCTTAAGCCTCAATAAAGCTTGC-3′), and RNase-free water. Reactions were performed on an Azure Cielo Real-Time PCR System using the following cycling conditions: 95°C for 1 min, followed by 45 cycles of 95°C for 10 s, 63.5°C for 1 min, and 72°C for 10 s. This approach enables sensitive detection of extracellular viral RNA and provides a direct measure of viral output over time. At defined time points, cells were harvested for downstream analyses.

### Analysis of cell-associated HIV-1 RNA

At the indicated time points, cell-associated nucleic acids were isolated from infected macrophages using the QIAGEN AllPrep DNA/RNA Mini Kit according to the manufacturer’s instructions. Briefly, macrophages were lysed directly in the culture wells using the provided lysis buffer, allowing for the simultaneous purification of both genomic DNA and total RNA from the same sample. Lysates were homogenized and processed through spin-column–based purification steps to separately isolate DNA and RNA fractions. Purified RNA was eluted in nuclease-free water. Nucleic acids were quantified prior to downstream analyses and subsequently used for quantitative PCR (qPCR) assays to evaluate cell-associated HIV RNA levels in infected macrophages. Cell-associated HIV-1 RNA was quantified by RT-qPCR using Volcano 2G DNA Polymerase (Medix Biochemica; Cat. No. 8100M). Reactions were performed in a final volume of 20 μL containing 4 μL of RNA template, 5× Volcano buffer, dNTPs, anti-Taq antibody, C1/f forward primer (5′-TAGACCAGATCTGAGCCTGGGA-3′), DnMatch3/r reverse primer (5′-ACAACAGACGGGCACACACTA-3′), 2n4nrB probe (5′-CCACTGCTTAAGCCTCAATAAAGCTTGC-3′), and RNase-free water. Amplification was performed on an Azure Cielo Real-Time PCR System using the following cycling conditions: 95°C for 1 min, followed by 44 cycles of 95°C for 5 s, 62°C for 15 s, and 72°C for 15 s. This approach targets the LTR region to amplify a region from U5 to the start of R.

### Fluorescence imaging

Imaging-based assessment of cell viability and HIV-1 infection was performed by immunofluorescence staining for viability, HIV-1 Gag, and nuclear content using DAPI. 50,000 MDM were seeded in black 96-well plates (Corning/Costar), infected with VSV-G pseudotyped HIV-1NL4-3 and treated with compounds. All subsequent steps were performed at room temperature. Cells were stained with 1 mM YOYO-1 iodide (Invitrogen) for 10 min, washed with DPBS (Gibco), and subsequently fixed with 4% paraformaldehyde (PFA; Cytek) for 20 min. Following fixation, cells were permeabilized for 15 min using 0.1% Triton X-100 (Sigma) and blocked for 30 min with 0.5% bovine serum albumin (BSA; Sigma) and 0.5% normal goat serum (Jackson ImmunoResearch) in DPBS to reduce nonspecific antibody binding. HIV-1 infection was detected using a mouse anti-Gag-PE antibody (KC-57; Beckman Coulter), followed by a secondary goat anti-mouse IgG-AF647 (Polyclonal; Invitrogen). Nuclei were counterstained with DAPI (BioLegend). Plates were washed with DPBS prior to imaging. Fluorescent images were acquired on the ImageXpress Pico (Molecular Devices) using appropriate filter cubes for YOYO-1, AF647, and DAPI. The percentage of Gag^+^ cells was determined using CellReporterXpress software (Molecular Devices).

### Live Imaging

50,000 MDM were seeded in black 96-well plates (Corning/Costar), infected with VSV-G pseudotyped HIV-1NL4-3 containing a GFP reporter gene, and treated with compounds for 16hrs whereafter 1.5 mg/mL Propidium Iodide (PI; Biolegend) was added to each well. Fluorescent images were acquired on the ImageXpress Pico (Molecular Devices) at 16h, 20h and 24h using appropriate filter cubes for GFP and PI.

### Statistical Analysis

All statistical analyses were performed using GraphPad Prism version 11.0.2 (GraphPad Software). Data are presented as mean ± SD for individual donor experiments and mean ± SEM for pooled data from independent donors. A *P* value of <0.05 was considered statistically significant. For Figure 1, differences in cell viability among treatment groups at each drug concentration were analyzed using an ordinary two-way ANOVA followed by Dunnett’s multiple-comparisons test, with each treatment compared to the corresponding DMSO control. For Figure 2, dose-response curves were generated by nonlinear regression using a variable-slope (four-parameter logistic) model. Infection data were normalized to the corresponding DMSO-treated control for each donor, and the half-maximal effective concentration (EC₅₀) was calculated with 95% confidence intervals. For the pooled extracellular HIV-1 RNA and cell-associated HIV-1 RNA datasets (Figures 3C and 4C), data were first normalized to the corresponding DMSO-treated control within each donor and expressed as fold change over DMSO. Statistical comparisons were performed using an ordinary two-way ANOVA followed by Tukey’s multiple-comparisons test to compare all treatment groups at each time point. Exact *P* values from these pairwise comparisons are displayed in the heat maps (Figures 3D and 4D). For Figure 5A, differences in the percentage of HIV-1 Gag-positive cells among treatment groups were analyzed using an ordinary one-way ANOVA followed by Dunnett’s multiple-comparisons test, with each treatment compared to the DMSO control.

## ACKNOWLEDGMENTS

Supported in part by a research grant from Investigator-Initiated Studies Program of Merck Sharp & Dohme LLC. The opinions expressed in this paper are those of the authors and do not necessarily represent those of Merck Sharp & Dohme LLC. This research was also supported by the Miami Center for AIDS Research (CFAR) at the University of Miami Miller School of Medicine of the National Institutes of Health under award number P30AI1073961, which is supported by the following NIH Co-Funding and Participating Institutes and Centers: NIAID, NCI, NICHD, NHLBI, NIDA, NIMHD, NIA, NIDDK, NIDCR, NIMH, NINR, NIGMS, and OD as well as grants R01MH116701, R01DK131533, R01MH128022-01, R01DA058437 from the National Institutes of Health.

